# Wild type domestication: loss of intrinsic metabolic traits concealed by culture in rich media

**DOI:** 10.1101/2024.09.12.612796

**Authors:** Ben Vezina, Helena B. Cooper, Jessica A. Wisniewski, Matthew H. Parker, Adam W. J. Jenney, Kathryn E. Holt, Kelly L. Wyres

**Author notes:** Corresponding authors Ben Vezina; Helena B. Cooper; Kelly L. Wyres. These authors contributed equally (Ben Vezina, Helena B. Cooper).

## Abstract

**Background:** Bacteria are typically isolated on rich media to maximise isolation success, removing them from their native evolutionary context. This eliminates selection pressures, enabling otherwise deleterious genomic events to accumulate. Here we present a cautionary tale of these ‘quiet mutations’ which can persist unnoticed in bacterial culture lines.

**Methods:** We used a combination of microbiological culture (standard and minimal media conditions), whole genome sequencing and metabolic modelling to investigate putative *Klebsiella pneumoniae* L-histidine auxotrophs. Additionally, we used genome-scale metabolic modelling to predict auxotrophies among completed public genomes (n=2,637).

**Results:** Two sub-populations were identified within a *K. pneumoniae* frozen stock, differing in their ability to grow in the absence of L-histidine. These sub-populations were the same ‘strain’, separated by eight single nucleotide variants and an insertion sequence-mediated deletion of the L-histidine biosynthetic operon. The His^-^ sub-population remained undetected for >10 years despite its in inclusion in independent laboratory experiments. Genome-scale metabolic models predicted 0.8% public genomes contained ≥1 auxotrophy, with purine/pyrimidine biosynthesis and amino acid metabolism most frequently implicated.

**Discussion:** We provide a definitive example of the role of standard rich media culture conditions in obscuring biologically relevant mutations i.e. nutrient auxotrophies, and estimate the prevalence of such auxotrophies using public genome collections. While the prevalence is low, it is not insignificant given the thousands of *K. pneumoniae* that are isolated for global surveillance and research studies each year. Our data serve as a pertinent reminder that rich-media culturing can cause unnoticed wild type domestication.

## Introduction

Whole genome sequencing of environmental and clinical microbes is crucial for understanding microbial diversity and distribution (1, 2), predicting antimicrobial resistance and tracking outbreaks (3). Prior to sequencing, isolates are enriched using nutrient-dense growth media like Luria-Bertani (LB) to maximise isolation success in large-scale studies (1, 2). However, what often goes unacknowledged is that isolation and sequencing represent a snapshot of an organism. Genomes and populations are in constant flux, guided by evolutionary selection pressures such as competition for space (4) and nutrients (5), bacteriophage predation (6) and antibiotics (7). Growth in rich media removes these pressures, permitting deleterious genomic events to accumulate. In this study, we present a cautionary tale of these ‘quiet mutations’ and how they can persist without notice. We also estimate the frequency of these events across public *Klebsiella pneumoniae* Species Complex (*Kp*SC) genomes.

## Methods

### Bacterial strains and culture

Two *Kp*SC isolates collected from hospitalised patients in Melbourne, Australia were used in this study: *K. pneumoniae* INF018 and *K. variicola subsp. variicola* INF232, both causing urinary tract infections (3). INF018 was part of a reported transmission cluster (8) and was used in two subsequent studies focusing on metabolism (3, 9) and one exploring passage through an *in vitro* bladder model (10), from which additional genome sequences were generated (**Figure 1A**). Isolates were grown as previously reported (11). Briefly, 5 mL cultures were grown at 37°C, shaking at 200 RPM overnight in broth: M9 minimal media (Sigma) containing either 20mM D-glucose, 20mM L-histidine, or both. Isolates were plated onto LB or M9 plates containing the same substrates.

**Figure 1.**
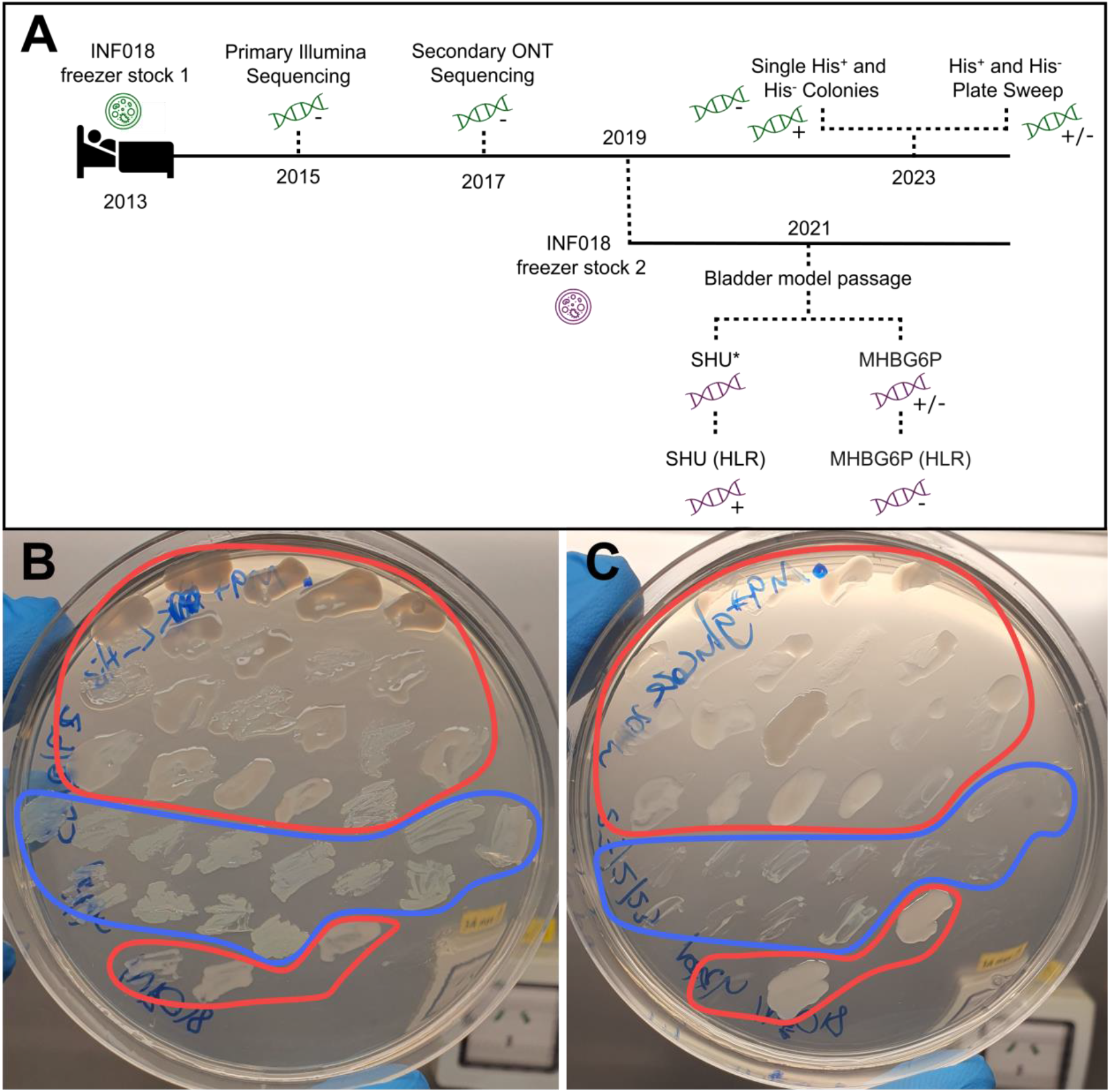
**A:** Timeline of *K. pneumoniae* INF018, with dark lines representing different frozen stocks (green and purple) and dotted lines showing independent subculturing events. Sequencing is represented with a DNA icon coloured by the frozen stock they originated from, with a “+” indicating a His^-^colony, “-” indicating a His^-^ colony and “+/-” indicating mixed colonies. Sequencing for the *in vitro* bladder model was done using INF018 passaged in either cation-adjusted Mueller–Hinton broth (MHBG6P) or synthetic human urine (SHU), including high-level Fosfomycin-resistant (HLR) subpopulations from each media (10). Sequencing data for INF018 SHU (labelled with an asterisk) is unavailable as the genome failed quality control (10). **B & C:** Patch plates showing heterogeneous population of His^+^ and His^-^ sub-populations in clinical frozen stocks. Colonies within the red boundaries show *K. pneumoniae* INF018 whereas the blue boundary show *K. variicola* subsp. *variicola* INF232. Colonies match positions on each plate. **B:** M9 agar + 20 mM L-histidine. **C:** M9 agar + 20 mM D-glucose.

### DNA isolation and sequencing

Overnight cultures were pelleted at 8000x g and DNA extracted using GenFind V3 (Beckman Coulter). DNA libraries were constructed using Illumina DNA Library Prep Kit, (M) Tagmentaton (Illumina) and sequenced on the Illumina NovaSeq 6000 at 2×250bp. Reads and assemblies were deposited at Genbank (BioProject PRJNA1115910, further details at Figshare (doi:10.6084/m9.figshare.25864864).

### Genomic analysis

Genomes were assembled using Unicycler v0.4.7 with the --keep 0 option (12) and annotated with Bakta v1.8.1 (13) using the ‘--gram –’ option and v1.8.1 database. Kaptive 3 (14-16) was used for determining K/O locus intactness. Read and contig mapping to a completed genome representing a His^+^ isolate from the same transmission cluster (accession: GCA_904863215.1) was performed with minimap2 v2.26 (17), using SAMtools v1.19 (18) for depth estimation. Genome-wide single nucleotide variant (SNV) distances were calculated via read mapping and variant calling against the completed INF018 genome (accession: SAMEA3356961, His^-^) using RedDog v1beta.11 (https://github.com/katholt/RedDog). ISEScan v1.7.2.3 (19) was used to identify Insertion Sequences.

Completed public *Klebsiella* genomes (n=2,637) were obtained from Genbank, accessed: 29/01/2024 (doi:10.6084/m9.figshare.25864864). Bactabolize (20) v1.0.2 was used to construct metabolic models (using the *Kp*SC pan v2.0.2 reference) and predict growth phenotypes (11).

## Results

### Heterogeneous sub-population in clinical isolate sample

We previously reported metabolic models (11) which unexpectedly predicted L-histidine auxotrophies for two isolates (*K. pneumoniae* INF018 and *K. variicola* INF232). L-histidine biosynthesis is essential for cellular growth and protein production and is conserved across all bacteria (21). Both genomes were complete (3), ruling out misassembly as an explanation, and both isolates were subsequently tested *in vitro* to confirm L-histidine auxotrophy (11). In parallel, another team member, unaware of these data, performed an independent experiment using the same INF018 frozen stock, which demonstrated growth in M9 + D-glucose without L-histidine.

When these inconsistencies were realised, the original frozen stocks were plated onto LB and colonies patched onto M9 + D-glucose and M9 + L-histidine (**Figure 1B & C**). All colonies grew on L-histidine, and 19/23 INF018 colonies grew on D-glucose whereas 0/13 INF232 colonies were able to grow on L-histidine. This confirmed the INF018 frozen stock was a heterogeneous population, consisting of His^+^ and His^-^ sub-populations, while INF232 was entirely His^-^. Compared to His^+^, His^-^ mutants displayed a flatter, paler colony morphology.

### Genomics confirmed deletion of L-histidine biosynthetic operon

Single INF018 His^+^ and His^-^ colonies were whole-genome sequenced to confirm the heterogeneous population (**Figure 1**). Compared to the complete INF018 genome (3), the INF018 His^+^ genome had eight SNVs while INF018 His^-^ had zero, indicating they were the same strain (22). Gorrie *et al*. had previously reported the INF018 frozen stock contained an ESBL plasmid (*bla*_CTX-M-15_), which was lost during culture and original sequencing (3), but we recovered the plasmid in both His^+^ and His^-^ colony genomes.

Read mapping indicated the INF018 His^-^ sub-population had a 19,388 bp deletion spanning the L-histidine biosynthetic operon (*his*) and part of the adjacent O locus (**Figure 2**). The loss of O-antigen would prevent K-antigen anchoring, which may have contributed to the colony morphology variation (**Figure 1B & C**). The chromosomal deletion event was likely mediated by insertion of IS5_222 introduced into the genome on the ESBL plasmid, and found exclusively on plasmids in closely related His^+^ *K. pneumoniae* strains (3).

**Figure 2:**
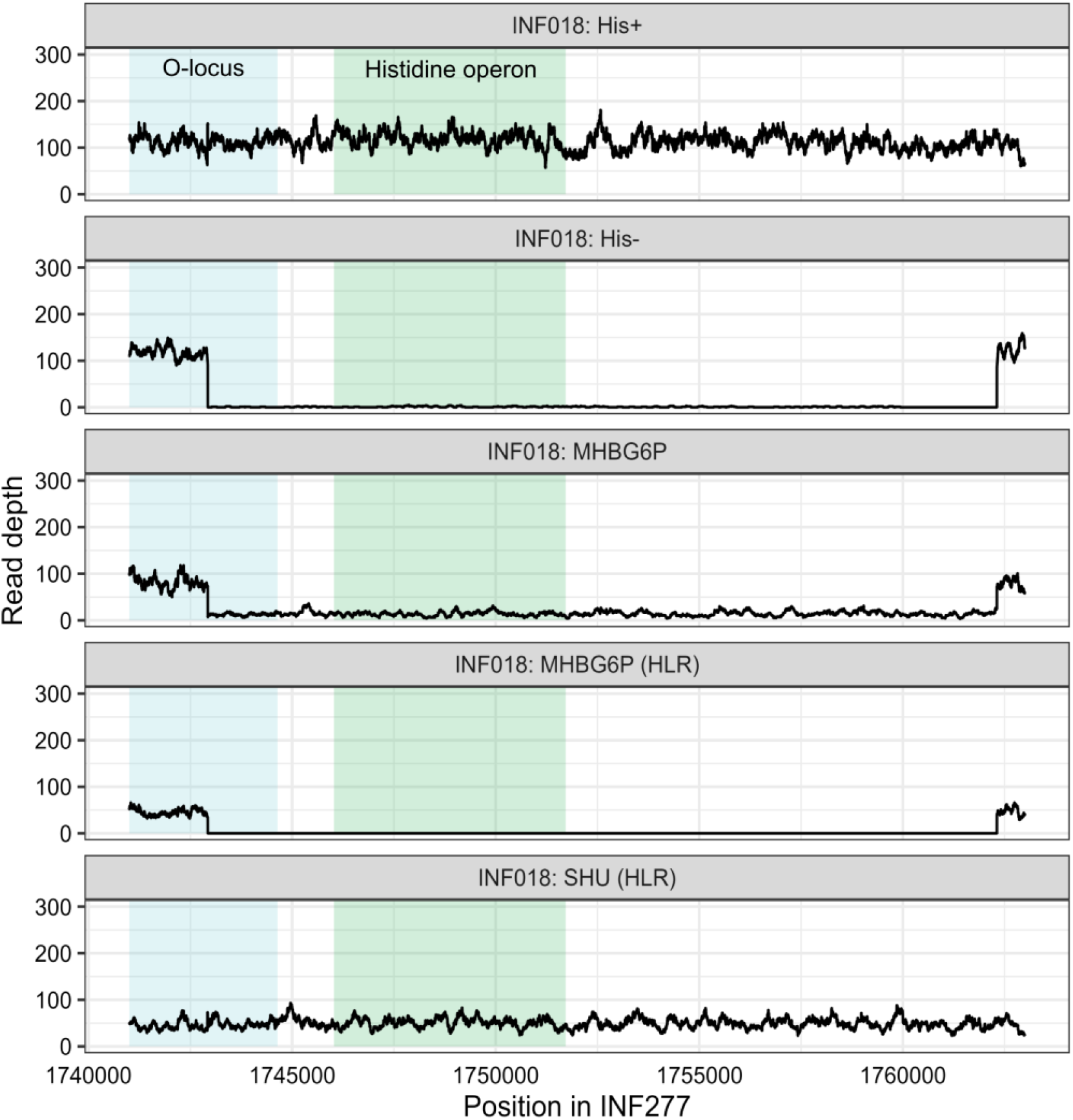
Read mapping of experimentally verified INF018 His^+^ and His^-^ strains as well as INF018 reads from the *in vitro* bladder model experiment (accessions: SAMN17846711, SAMN17846713 & SAMN21465650) (10) wherein INF018 was passaged in cation-adjusted Mueller–Hinton broth (MHBG6P) or synthetic human urine (SHU), prior to sequencing. High-level Fosfomycin-resistant (HLR) subpopulations from each passage were also sequenced (10). Reads were aligned to *K. pneumoniae* INF277 (accession: SAMN06112222, His+), which is part of the same transmission cluster as INF018 (3). The green box highlights the location of the L-histidine biosynthesis operon, and the blue box highlights the 3’ region of the O-antigen locus.

Prior to this work, INF018 was used in an *in vitro* bladder model experiment (10) where it was passaged in cation-adjusted Mueller–Hinton broth (MHBG6P) or synthetic human urine (SHU) with/without fosfomycin and subjected to *de novo* whole-genome sequencing (**Figure 1A)**. Re-analysis suggests that a His^+^ isolate was used for SHU experiments while a His^-^ isolate was used for MHBG6P, suggesting that the second INF018 freezer stock also contained a mixed population (**Figure 1A & 2**). However, there is some read mapping to the *his* operon in the MHBG6P 24-hour passaged genome that was absent in the 48-hour passaged genome, implying a gradual loss in the operon overtime (**Figure 2**).

### Estimating auxotrophies from public genome data

Given that we had identified two L-histidine auxotrophs among a collection of just 507 isolates (0.4%) (11), we sought to estimate the general prevalence of such auxotrophs among public genome collections. We constructed metabolic models for all complete (20) *Kp*SC genomes in Genbank (n=2,637). Twenty (0.8%) were predicted as auxotrophs on M9 + D-glucose media, with the most frequent auxotrophies being purine and pyrimidine biosynthesis and amino acid metabolism (**Figure 3**).

**Figure 3:**
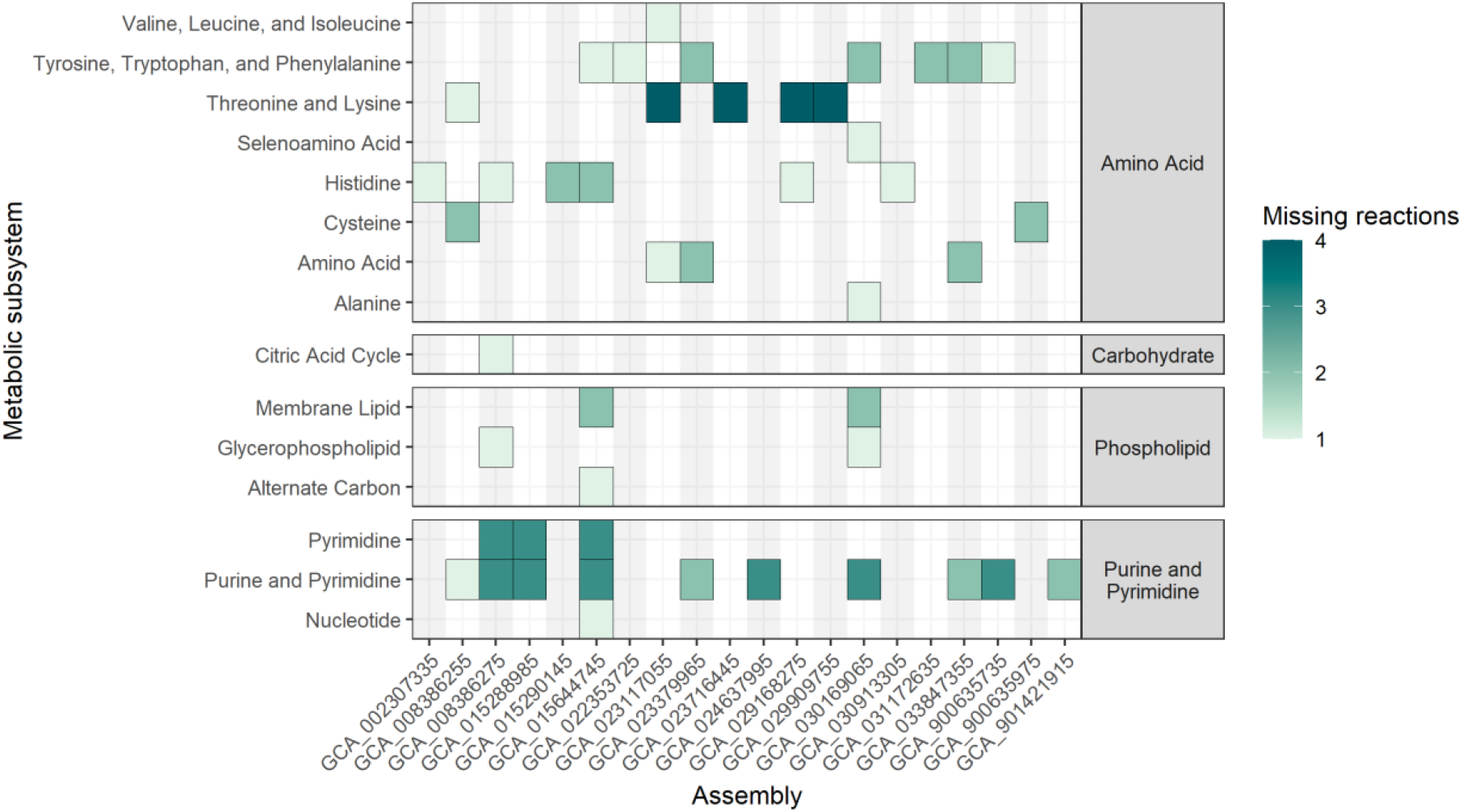
Metabolism auxotrophies identified in the 20 putatively auxotrophic isolates. Counts of individual missing reactions are shown by colour. Auxotrophs are split by the subsystem and general metabolic class. Median missing reactions resulting in substrate auxotrophism was 3 (range 1 to 25).

## Discussion

Loss of plasmids during culturing (3, 25, 26) and spontaneous mutants with varying colony morphology (27, 28) have been described previously, as have passage-induced mutations in historical and globally-distributed laboratory strains (23, 24). For example, passage of *Burkholderia pseudomallei* strain K96243 in LB broth for 28 days impacted virulence, protein expression and regulation of metabolism (24). However, to our knowledge there are few discussions of spontaneous nutrient auxotrophs occurring over the timescale of standard clinical microbiological diagnostic protocols and/or overnight cultures for DNA extractions. Current recommendations for *Klebsiella* isolation include growth in LB + ampicillin then Simmons Citrate + Inositol (29) or MacConkey/CHROMagar/horse blood agar (30). The two isolates used in this study (*K. pneumoniae* INF018 and *K. variicola* INF232) were originally collected from patient samples plated onto MacConkey then grown in LB broth prior to sequencing (3). These rich medias would support the within-isolate heterogeneity we observed, eventually resulting in the complete loss of His^+^ sub-populations. However, it is also possible that both sub-populations were present in the patient (urine is histidine rich), but in either case the use of rich media enabled the His^-^ population to persist and outcompete His^+^ populations unnoticed (10). We therefore propose that these spontaneous auxotrophies act like *‘*quiet mutations’ - a phenotypic parallel to silent mutations. These occur when DNA changes cause a phenotype alteration that remains unnoticed in standard culture conditions.

Our analysis predicted at least 0.8% of completed, high-quality *Kp*SC genomes represent isolates with ≥1 auxotrophy (**Figure 3**), indicating that the phenomenon is rare but not insignificant considering the scale of modern genomic sequencing i.e. 100,000 *K. pneumoniae* genomes are now available in NCBI Pathogens. This may further be an underestimation of quiet mutations considering that we focused solely on metabolic gene content but expect that non-metabolic gene content would also be impacted.

This cautionary tale has served as a good self-reminder when studying microbial ecology to make sure we are observing the wild type.

## Notes

### Competing Interest Statement

The authors have declared no competing interest.

